# Neutralization of SARS-CoV-2 VOC 501Y.V2 by human antisera elicited by both inactivated BBIBP-CorV and recombinant dimeric RBD ZF2001 vaccines

**DOI:** 10.1101/2021.02.01.429069

**Authors:** Baoying Huang, Lianpan Dai, Hui Wang, Zhongyu Hu, Xiaoming Yang, Wenjie Tan, George F. Gao

## Abstract

Recently, the emerged and rapidly spreading SARS-CoV-2 variant of concern (VOC) 501Y.V2 with 10 amino acids in spike protein were found to escape host immunity induced by infection or vaccination. Global concerns have been raised for its potential to affect vaccine efficacy. Here, we evaluated the neutralization activities of two vaccines developed in China against 501Y.V2. One is licensed inactivated vaccine BBIBP-CorV and the other one is recombinant dimeric receptor-binding domain (RBD) vaccine ZF2001. Encouragingly, both vaccines largely preserved neutralizing titres, with slightly reduction, against 501Y.V2 authentic virus compare to their titres against both original SARS-CoV-2 and the currently circulating D614G virus. These data indicated that 501Y.V2 variant will not escape the immunity induced by vaccines targeting whole virus or RBD.

## Maintext

The rollout of vaccines is the hope to control the coronavirus disease 2019 (COVID-19) pandemic and reboot economy and society(*1*). However, the recent emergence of severe acute respiratory syndrome coronavirus-2 (SARS-CoV-2) variants raised global concerns because of their enhanced transmission and potential viral escape of host immunity elicited by natural infection or vaccination. Variants containing D614G mutation in spike (S) protein was first reported from the middle of 2020, which significantly increased the transmission rate and became dominant in circulating strains ever since(*2*). Evolved from D614G mutants, recent circulating isolates from Republic of South Africa (501Y.V2 variant) introduced further mutations that escape neutralization by COVID-19 convalescent plasma and sera from human receiving licensed mRNA vaccines expressing SARS-CoV-2 S protein(*3-5*). It also dramatically decreased the protective efficacy for trimeric S protein-based vaccine in phase 3 clinical trial in South Africa(*6*).

501Y.V2 variant emerged in more and more countries, and was first isolated in China on January 6, 2021 from an airline pilot of South African nationality(*7*). This variant, GDPCC strain, contains 10 amino acid mutations sites in S protein corresponded to the features of the variant of concern (VOC) South African 501Y.V2, with 5 (D80A, ΔL242, ΔA243, ΔL244 and R246I) located at N-terminal domain (NTD), 3 (K417N, E484K and N501Y) in receptor-binding domain (RBD), and the other two in CTD2 domain and S1/S2-S2’ region (Fig.1).

**Fig 1.**
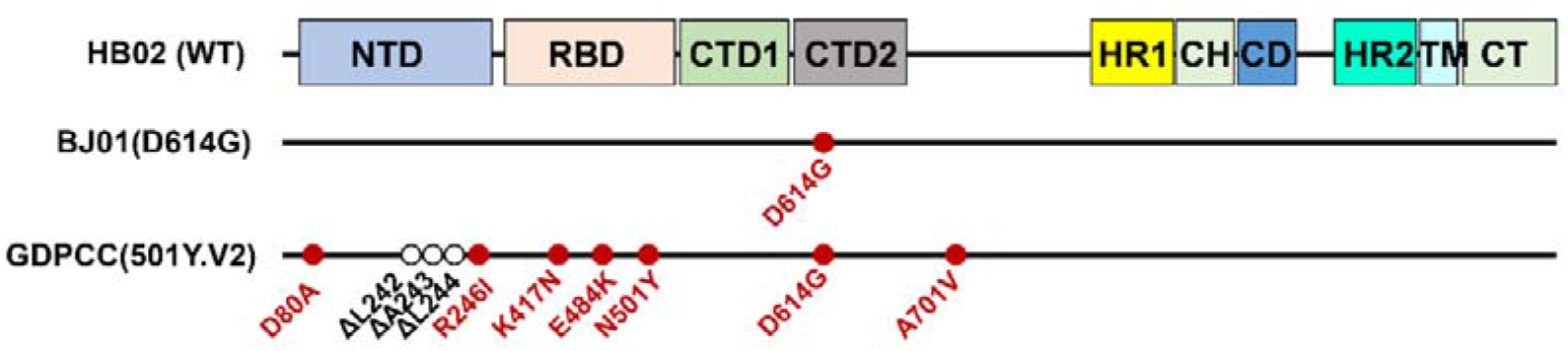
Schematic demonstration of the spike protein of SARS-CoV-2 HB02 (WT), BJ01 (D614G) and GDPCC (501Y.V2). The mutation sites were denoted and marked as cycles, with the point mutation colored in red and the deletion colored in white.

The effectiveness of current vaccines against this VOC is of high importance to guide the ongoing vaccination program worldwide. To answer this question, we evaluated two representative COVID-19 vaccines developed in China for their neutralizing activities against 501Y.V2 variant. One is BBIBP-CorV, a licensed COVID-19 inactivated vaccine(*8, 9*). The other one is ZF2001, a protein subunit vaccine targeting S protein RBD currently in phase 3 clinical trials(*10, 11*). Both vaccines showed good immunogenic in their clinical trials(*9, 11*). For instance, ZF2001 induced neutralizing GMTs two times greater than that from convalescent samples(*11*). We chose 12 serum samples for each vaccine from clinical trial participants covering a range of different neutralizing titers (Table 1). The neutralizing activities of these serum samples against live SARS-CoV-2 strains GDPCC (501Y.V2) were measured by microcytopathogenic effect assay. SARS-CoV-2 strains HB02 (WT) (*12, 13*) and BJ01 (D614G) (*14*) were tested as the control. Impressively, all 12 serum samples from either ZF2001 or BBIBP-CorV recipients largely preserved neutralization of 501Y.V2 variant, with slightly reduced geometric mean titres (GMTs) compared with their titres against WT or D614G strain (Fig. 2 and Table 1). For ZF2001, the GMTs declined for 1.6-fold from 106.1 (95% CI, 75.0-150.1) to 66.6 (95% CI, 51.0-86.9) (Fig.2A). While for BBIBP-CorV, the decline is also 1.6-fold, GMTs from 110.9(95% CI, 76.7-160.2) to 70.9(95% CI, 50.8-98.8) (Fig.2B). These reductions are significantly less than those reported previously for convalescent plasma (more than 10-folds)(*4*) or antisera from mRNA vaccine recipients (more than 6-folds) (*3, 5*).

**Table 1:**
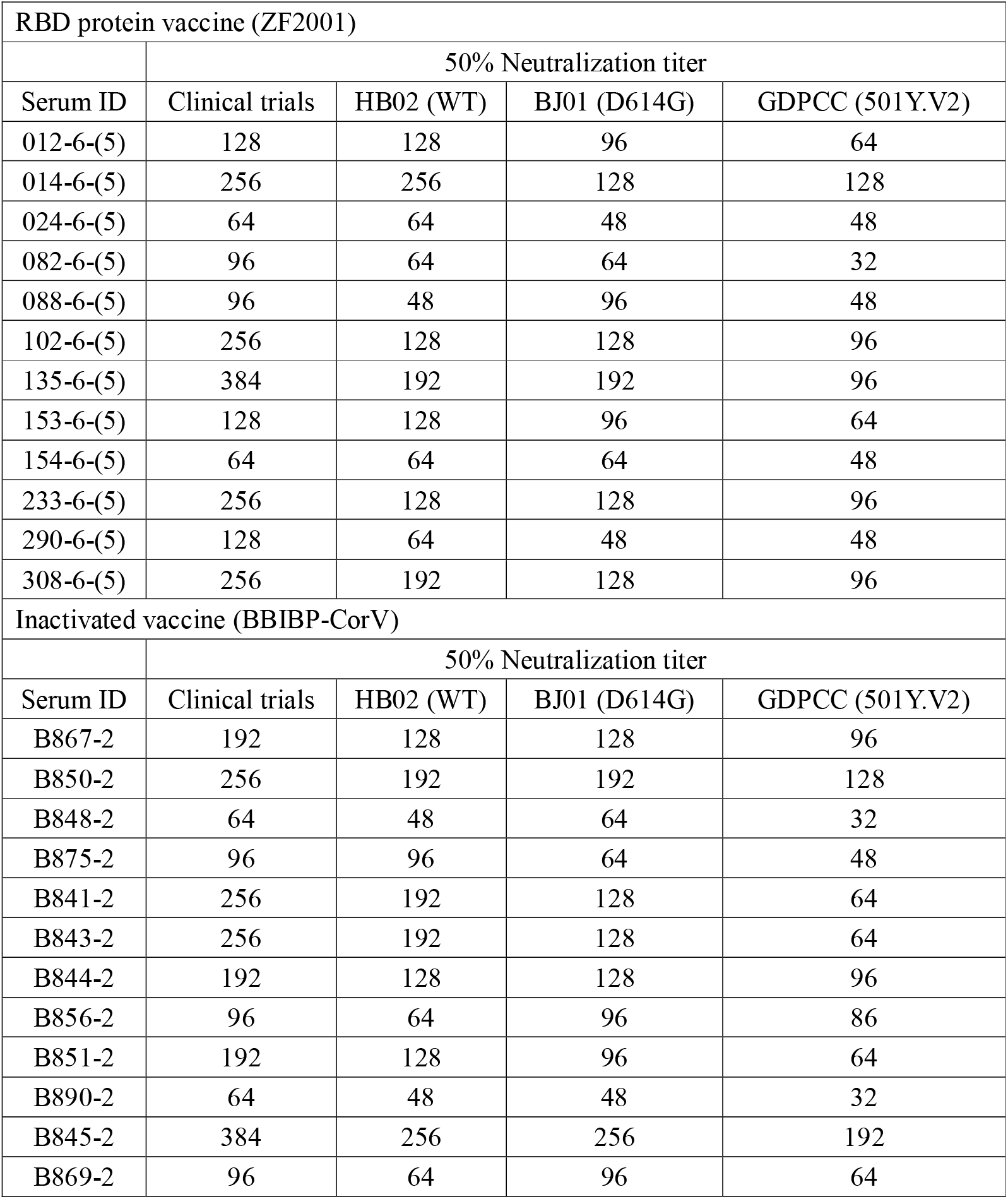
Information for serum samples tested in this study.

| RBD protein vaccine (ZF2001) |  |  |  |  |
| --- | --- | --- | --- | --- |
|  | 50% Neutralization titer |  |  |  |
| Serum ID | Clinical trials | HB02 (WT) | BJ01 (D614G) | GDPCC (501Y.V2) |
| 012-6-(5) | 128 | 128 | 96 | 64 |
| 014-6-(5) | 256 | 256 | 128 | 128 |
| 024-6-(5) | 64 | 64 | 48 | 48 |
| 082-6-(5) | 96 | 64 | 64 | 32 |
| 088-6-(5) | 96 | 48 | 96 | 48 |
| 102-6-(5) | 256 | 128 | 128 | 96 |
| 135-6-(5) | 384 | 192 | 192 | 96 |
| 153-6-(5) | 128 | 128 | 96 | 64 |
| 154-6-(5) | 64 | 64 | 64 | 48 |
| 233-6-(5) | 256 | 128 | 128 | 96 |
| 290-6-(5) | 128 | 64 | 48 | 48 |
| 308-6-(5) | 256 | 192 | 128 | 96 |
| Inactivated vaccine (BBIBP-CorV) |  |  |  |  |
|  | 50% Neutralization titer |  |  |  |
| Serum ID | Clinical trials | HB02 (WT) | BJ01 (D614G) | GDPCC (501Y.V2) |
| B867-2 | 192 | 128 | 128 | 96 |
| B850-2 | 256 | 192 | 192 | 128 |
| B848-2 | 64 | 48 | 64 | 32 |
| B875-2 | 96 | 96 | 64 | 48 |
| B841-2 | 256 | 192 | 128 | 64 |
| B843-2 | 256 | 192 | 128 | 64 |
| B844-2 | 192 | 128 | 128 | 96 |
| B856-2 | 96 | 64 | 96 | 86 |
| B851-2 | 192 | 128 | 96 | 64 |
| B890-2 | 64 | 48 | 48 | 32 |
| B845-2 | 384 | 256 | 256 | 192 |
| B869-2 | 96 | 64 | 96 | 64 |

**Fig 2.**
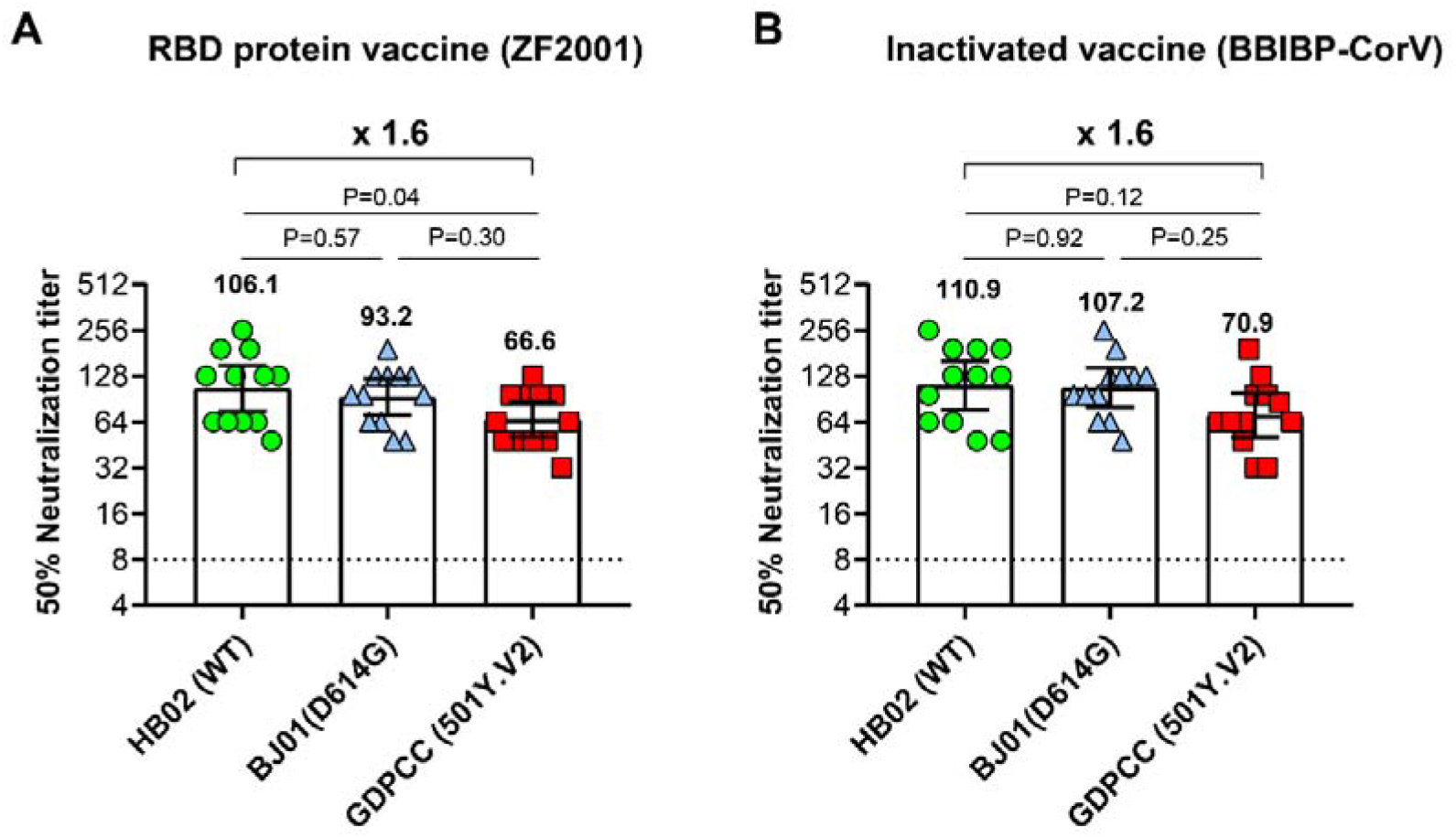
Neutralization titres of 24 antisera from vaccine BBIBP-CorV or ZF2001 recipients against authentic SARS-CoV-2 and its variants, D614G and 501Y.V2. (A-B) N=12 representative antisera each from ZF2001 (A) and BBIBP-CorV (B) vaccine recipients were tested for their neutralizing activity to authentic SARS-CoV-2 HB02 (WT), BJ01 (D614G) and GDPCC (501Y.V2) via microcytopathogenic effect assay. Shown are the geometric mean with 95% CI. P values were analyzed with One-way ANOVA test.

These results suggest that the 501Y.V2 variant does not escape the immunity induced by vaccines targeting S protein RBD (ZF2001) or whole virus (BBIBP-CorV). The potential 1.6-fold reduction of neutralizing GMTs should be taken into account for its impact for the clinical efficacy of these vaccines. For both vaccines, antisera neutralize both variant 501Y.V2 and D614G, the one currently circulating globally, without statistical significances. For ZF2001, a slight significance (P=0.04) between variant 501Y.V2 and the HB02 may due to the sample selection and size. For the neutralization-reduction discrepancy between our protein-based vaccine and mRNA vaccine needs further investigation in the future.

## Acknowledgements

This work was supported by the National Program on Key Research Project of China (2020YFA0907101, 2016YFD0500301), the National Natural Science Foundation of China (NSFC) (82041041, 82061138008), National Mega Projects of China for Major Infectious Diseases (2016ZX10004001-003). Lianpan Dai is supported by Youth Innovation Promotion Association of the CAS (2018113).

## Supplementary Materials for

### Material and methods

#### Viruses and titration

The SARS-CoV-2 virus 19nCoV-CDC-Tan-HB02 (short as HB02), 19nCoV-CDC-Tan-BJ01 (short as BJ01) and 19nCoV-CDC-Tan-GDPCC (short as GDPCC) were used in our experiments*(1-4)*. Virus titer was determined by a micro-cytopathogenic efficiency (CPE) assay on Vero cells as described previously*(5)*.

#### Neutralization assay

The serum were inactivated in a 56□ water bath for 30 min, then successively diluted 1:4 to the required concentration by a 2-fold series. An equal volume of challenge virus solution containing 100 CCID_50_ virus was added. After neutralization in a 37□ incubator for 2 h, a 1.0∼2.5×10^5^/ml cell suspension was added to the wells and cultured in a CO_2_ incubator at 37□ for 4 days. Titers expressed as the reciprocal of the highest dilution protecting 50% cell from virus challenge. A neutralization antibody potency < 1:4 is negative, while that ≥1:4 is positive.

#### Serum samples from clinical trial participants

For the 12 Serum samples from BBIBP-CorV vaccination(*6, 7*), vaccine recipients received BBIBP-CorV containing 4 μg total protein on days 0 and 21, blood samples were taken from participants for serology tests 28 days after second vaccination, covering a range of different neutralizing titers, the ClinicalTrials.gov Identifier is NCT04510207.

For the 12 Serum samples from ZF2001 vaccination (*8*), vaccine recipients received ZF2001 containing 25 μg vaccine dose on days 0, 30, 60. Blood samples were taken from participants for serology tests 14 days after third vaccination, covering a range of different neutralizing titers, the ClinicalTrials.gov Identifier is NCT04466085.

